# *De novo* Sequencing and Whole Transcriptome Analysis for Biosynthesis Pathway of Diosgenin in *Pedalium murex* L.: A Medicinal herb

**DOI:** 10.1101/2023.11.16.567345

**Authors:** Parul Tyagi, Rajiv Ranjan

**Affiliations:** Department of Botany, Dayalbagh Educational Institute, Dayalbagh Agra 282005, India

**Keywords:** *Pedalium murex*, transcriptome, diosgenin biosynthesis, transcription factor, SSR, homology, bioinformatics tools

## Abstract

Transcriptome-based investigations of candidate genes, critical pathways, and gene regulation in non-model species have been transformed by high-throughput RNA sequencing in different studies. *Pedalium murex* L. is one of the most important annual medicinal herbs of the *Pedaliaceae* family. Because of its anti-inflammatory, antioxidant, antimicrobial, and anticancerous properties, *P. murex* is widely used in traditional medicine to treat gonorrhoea, leucorrhoea, urinary disorders, gastrointestinal tract disorders, cough, and asthma. Steroidal diosgenin is the major bioactive compound of *P. murex*. However, transcriptional technologies have yet to be used to study the steroid diosgenin biosynthetic pathway of this herb. In this study, we performed a whole transcriptomic analysis of the root, fruit, and leaf tissues of *P. murex* with three biological replicates and obtained ∼6.77 Gb of clean raw data. A total of 148871 unigenes were assembled with an average length of N50 and 1167 bp. Putative functions could be annotated to 75198 unigenes based on a BLASTX search against the NR, Uniprot, KEGG, Pfam, GO, and COG databases. Most of the unigenes related to steroidal diosgenin backbone biosynthesis were up- regulated in the root, fruit and leaf, except for the MVD gene in the leaf. qRT-PCR further verified the differential expression analysis of selected genes. It shows the highest homology with *Sesamum indicum, Handroanthus impetiginosus, Erythranthe guttata, Oleaeuropaea* var*. sylvestris,* and *Dorcoceras hygrometricum*. A total of 21026 unigenes of transcription factors were assembled into transcription factor families. A total of 8760 unigenes of SSR were assembled. The transcriptome data presented here will make it easier to study the functional genomics of steroidal diosgenin biosynthesis and to change the genes of *P. murex* to make it more diosgenin.

## 1 INTRODUCTION

*Pedalium murex* is one of the most valuable medicinal herbs, commonly known as bada gokhuru and large calatrop, and belongs to the *Pedaliaceae* family. It is natively and extensively grown in China, Pakistan, India, Sri Lanka, and tropical Africa. It is an annual, glabrous succulent herb primarily found in waste places (Srinivasan 1942; Nadakarni 1982; Shukla and Thakur 1983; Bhakuni et al., 1992). *P. murex* is primarily used to treat sexual dysfunctions, but it has also been used to treat other conditions such as nocturnal emission, gonorrhoea, leucorrhoea, urinary disorders, gastrointestinal tract disorders, cough, and asthma. It also serves as a good gargle for mouth troubles and painful gums.

Among various bioactive chemical compounds, steroidal saponins (triterpenoids in nature) are the major group of secondary metabolites, which have received the most interest due to their nephroprotective, anti-inflammatory, antioxidant, antimicrobial, anticancerous, etc., activities, diosgenin being one of them (Rajashekar et al., 2012). Furthermore, the part of diosgenin named aglycone is the primary precursor of steroidal drugs such as contraceptives, testosterone, progesterone, and glucocorticoids (Kevalia and Patel, 2011; Patel et al., 2012; Anandalakshmi et al., 2016). Nowadays, the isolation of triterpenoids is in much demand owing to their pharmaceutical and pharmacological efficacy (Gomes-Carneiro et al., 1998). *P. murex* has been used to derive a variety of secondary metabolites with various biological implications, including steroidal saponins, flavonoids, alkaloids, fatty acids, vitamins, tannins, phenolic compounds, and so on (Rao et al., 1999). *P. murex* represents an alternative source of diosgenin because of its higher yield, negligible production cost, and shorter life cycle.

The medicinal and therapeutic importance of the species is due to the occurrence of steroidal diosgenin, one of the most structurally diverse and extensively distributed secondary metabolites in plants (Perez-Labrada et al., 2011). The diverse nature, number, and linkage pattern of sugar moieties in the aglycone skeleton contributes to the broad range of biological and pharmacological functions of steroidal diosgenin (Chaudhary et al., 2016). Although much steroidal diosgenin has been identified in *P. murex* (Tyagi et al., 2021), globally, diosgenin is used as an anticancerous and anti-ageing agent, besides its use as a precursor for the preparation of many steroidal drugs. Interestingly, *P. murex* accumulates almost triple diosgenin content (∼6.0%) compared to other explored medicinal plant species, namely *Asparagus* spp., *Chlorophytum* spp., *Dioscorea* spp., and *Trigonella* spp. (Sood et al., 2016).

Steroidal diosgenin is mainly synthesized from squalene through isopentenyl diphosphate (IPP) and dimethylallyl diphosphate (DMAPP) via cytosolic mevalonate (MVA) and plastidial 2-C-methyl-D-erythritol 4-phosphate (MEP) pathways, respectively. The cyclization of 2, 3-oxidosqualene synthesized from squalene is the first diversifying step for the biosynthesis of steroidal and triterpenoid diosgenin catalyzed by oxidosqualene cyclases (OSCs). In plants, cycloartenol synthase and lanosterol synthase mediate the cyclization of 2, 3-oxidosqualene to synthesize steroidal diosgenin (Haralampidis et al., 2002; Abe et al., 1993; Xu et al., 2004).

Despite *P. murex* numerous medicinal properties, no published genome or transcriptome sequence data is available in public databases. Given the lack of genomic and transcriptomic information and genome complexity (polyploidy and huge genome size), elucidating the complicated steroidal saponin biosynthesis pathway (Wang et al., 2018) at the genome level would be extremely difficult. The intricacy of medicinal plant genomes, as well as the cost of sequencing and processing resources, makes whole-genome sequencing difficult. As a result, only a few medicinal plants have had their entire genome sequenced. Transcriptome-based research has been transformed by next-generation high-throughput mRNA sequencing (Saito et al., 2013; Muranaka et al., 2013). Large-scale transcriptome sequencing is now possible thanks to recent developments in sequencing technology, which allow for gene expression and functional genomic research. Plant gene function data, as well as insight into the production of active components and their control, may be obtained quickly and efficiently using whole transcriptome research (Wang et al., 2009).

So, this study provided genetic information about this herb by de novo assembly of the Illumina *HiSeq* 2000 RNA-Sequencing platform for studying diosgenin biosynthesis pathway genes. The short reads were assembled using specific tools such as sequence platforms, assembly tools, and special bioinformatics analysis tools. These tools were applied to generate a complex technology system for genome and transcriptome sequences. These techniques can also be applied to identify transcript sequence polymorphisms and novel trans-splicing and splice isoforms (Ansorge and Heidelberg, 1991; Schuster, 2008; Thomas et al., 2007; Ansorge, 2009). Deep sequencing data and bioinformatics analysis for the several essential genes of diosgenin biosynthesis pathways were assembled. Most of the critical genes that are related to the diosgenin biosynthesis pathway were identified.

In this study, we have analyzed full-length mRNA derived from pooled tissues of root, fruit, and leaf of *P. murex*, sequenced using the Illumina platform, and successfully assembled isoform transcriptomes generated by alternative splicing. Identification of diosgenin-related genes and their functions based on expressed genes in all parts of herbs that have not previously been described. This transcriptome data will be useful for future genome assembly in *P. murex*, besides its use for developing functionally relevant molecular marker resources to assist population genetics and conservation studies.

## 2. Material and Methods

### Plant materials

Samples of all parts (root, fruit, and leaf) of *P. murex* were collected in September from the herbal garden of Dayalbagh Educational Institute, Dayalbagh, Agra, and are shown in **(Figure 1)**. After cleaning with ultrapure water, all parts were collected separately. These are immediately frozen in liquid nitrogen and stored at −80 °C until use.

**Figure 1:**
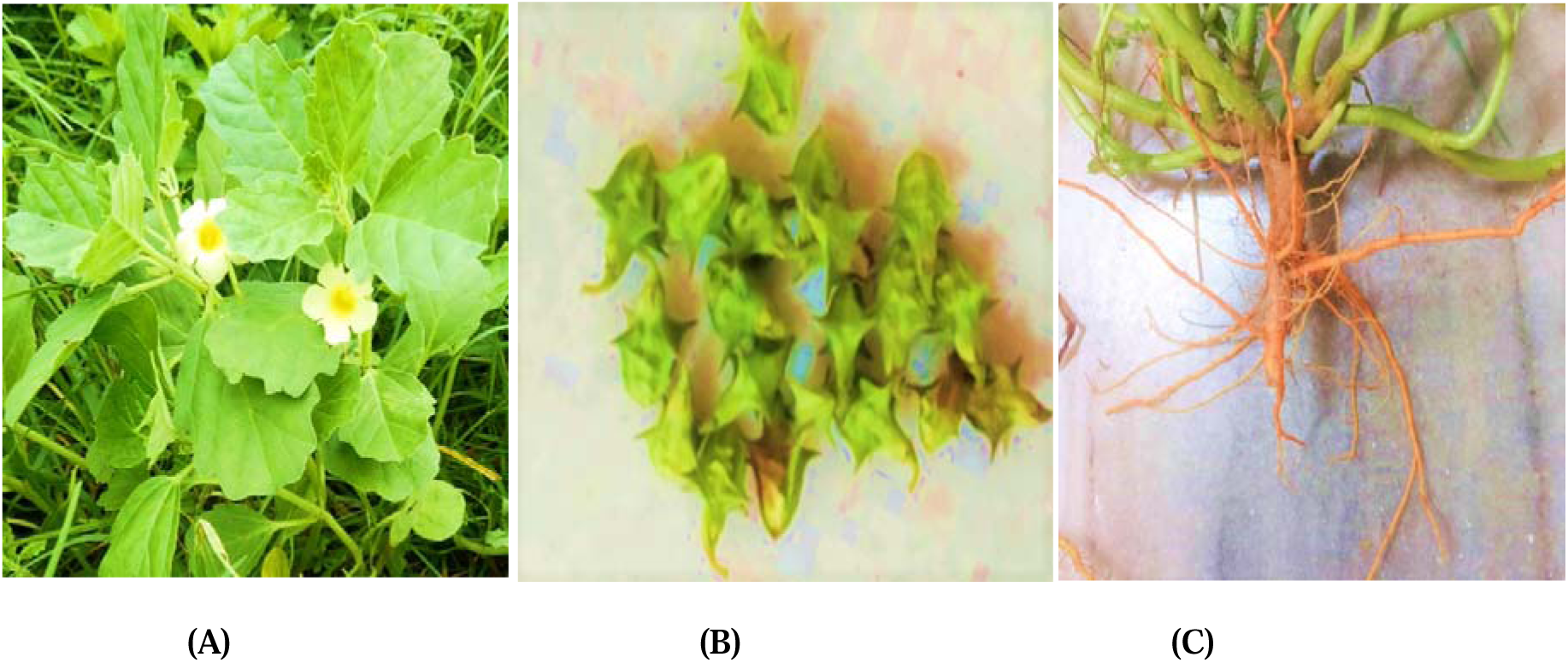
Description of leaf, fruit, and root tissues of *Pedalium murex*; (A) Leaf; (B) Fruit; (C) Root.

### Illumina RNA-Seq library Preparation and de novo *sequencing*

The total RNA from approximately 1.0 gm of each tissue of *P. murex* was extracted using the *RaFlex Total RNA isolation Kit* (Merck Millipore, Massachusetts, USA) as per the described protocol. Three replicates were employed for each experiment. *P. murex* root, fruit, and leaf tissue total RNA samples were pooled to form one equimolar (4 µl of each sample) concentration (Liu et al., 2013; Long et al., 2014; Liu et al., 2015; Ishihara et al., 2015). The quantity and quality of pooled samples were analyzed using a 1% formaldehyde agarose gel and a Nanodrop 8000 spectrophotometer (Thermo). For Illumina, mRNA was purified from the pooled samples. The pooled RNA samples were then subjected to cDNA library preparation using the *TruSeq Library Preparation Kit* (Illumina Inc., San Diego, CA, USA) as per its described protocol, with the higher RNA Integration Number (RIN) values (>9) being used for cDNA synthesis (Kim et al., 2019). The enriched mRNA was fragmented and converted into first-strand cDNA, followed by second-strand generation, A-tailing, adapter ligation, and finally ended by a limited number of PCR amplifications of the adaptor-ligated libraries. The amplified libraries were analyzed on the Bioanalyzer 2100 (Agilent Technologies) as per the manufacturer’s instructions. Library quantification and qualification were performed using the *DNA High Sensitivity Assay Kit* by Nanodrop 8000 spectrophotometer (Moraortiz et al., 2016). *P. murex cDNA* was sequenced by a paired-end (PE) 2×150 bp library on the Illumina *HiSeq* 2000 platform (Illumina, San Diego, California, USA) (Crawford et al., 2010).

### Functional annotation

All assembled unigenes were annotated by BLASTx analysis (Altschul et al., 1990) against the Nr (http://www.ncbi.nlm.nih.gov/), UniProt (http://www.uniprot.org/downloads), Pfam (http://pfam.xfam.org/), and Clusters of Orthologous Groups (COG) (http://www.ncbi.nlm.nih.gov/COG/) databases (Bairoch and Apweiler, 2000; Finn et al., 2014) with an E-value **<**1e-6. Only the top hit results were extracted for each unigenes of the transcriptome sample. Gene ontology (GO) (http://www.geneontology.org) terms were functionally classified based on Nr annotations by using *Blast2GO* (version 2.5.0; default parameter) (http://www.blast2go.de/) (Conesa et al., 2005). KEGG (http://www.genome.jp/kegg/) was used to draw metabolic maps. The KEGG analysis (Moriya et al., 2007; Kanehisa et al., 2014) results included KEGG Orthology (KO) numbers and enzyme commission (EC) numbers. Transcripts of transcription factors were aligned using BLASTX *Plant Transcription Factor Databases* (http://planttfdb.cbi.pku.edu.cn/).

### Analysis of genes encoding transcription factors (TFs)

The open reading frame (ORF) of each unigenes was discovered using the software getorf (EMBOSS:6.5.7.0) (Rice et al., 2000) to determine the transcription factor families represented in the *P. murex* transcriptome. The plant transcription factor database (PlnTFDB) was used to align these ORFs to all TF protein domains using BLASTX (e-value ≤ 1e-6) and the HMMER search technique (Mistry et al., 2013).

### Analysis of Quantitative Real-Time (qRT-PCR)

To validate RNA-*Seq* data through qRT-PCR analysis by taking the leaf, fruit, and root was conducted using 96 real-time PCR system (Bio-Rad, USA) (Agilent Aria (v1.5) Software) with a SYBR® Premix Ex Taq™ kit (Takara, China). Candidate reference qRT-PCR gene primers were designed using Primer Premier (version 5.0). Successive qRT-PCR experiments were done using triplicates, and melting curve analysis was performed following every amplification to confirm product specificity. To normalize, the alpha-tubulin gene (taken as control) from *P. murex* was used as an internal reference gene, and each sample was divided into three triplicates. The relative expression levels of each gene were calculated using the 2^−ΔΔCt^ method (Livak et al., 2001).

### SSR identification

Simple Sequence Repeats (SSRs) were found using the Micro Satellite software Perl script “MISA,” in which unigenes were utilized as reference material and transcript contigs were searched for SSRs (Moraortiz et al., 2016). The sequence was first made and mined for SSR motifs on both ends (dimer to hexamer nucleotide) with a length of 150 bp, and these were kept for di-, tri-, tetra-, and hexanucleotide repeats.

## 3. RESULTS

### Sequencing of RNA and *de novo* assembly

The increased applicability of NGS technologies, such as novel gene discovery, tissue-specific expression analysis, and the creation of sequence-based molecular marker resources, provides an excellent opportunity to dissect complex biosynthetic pathways and allows for a better understanding of the genomics of non-model plants (Unamba et al., 2015). To characterize the transcriptomics of *P. murex*nine cDNA libraries were considered that were prepared from the root, fruit, and leaf tissues to elucidate their whole expression patterns in *P. murex* by using the paired-end (PE) sequencing of the Illumina *Hiseq* 2000 platform. After removing adaptors, poly-A tails, primer sequences, short (150 bp) and low-quality sequences, a total of ∼6.77 GB of clean data was obtained. A total of 410 million raw reads were generated from pooled tissues of *P. murex*. A total of 16889795 duplicate reads were received. Furthermore, 75198 transcript scaffolds were assembled from all clean-reads. A total of 28.38% (26358323), 28.27% (26253769), 21.90% (20333352), and 22.45% (19918408) of ATGC composition percentages were obtained from all parts of *P. murex* **Table 1**. The high-quality reads and all isoforms were put together using the Trinity program (Grabherr et al., 2011) and the TGI clustering tool (TGICL) (Perteal et al., 2003) to get rid of duplicate sequences.

**Table 1:** Transcriptome data of whole herbs parts (root, fruit, and leaf) of *P. murex*.

| Samples of (root, fruit, and leaf tissues) | Raw data of *P. murex* |
| --- | --- |
| Total reads | 41091664 |
| Total duplicate reads | 16889795 (41.10%) |
| Total bases | 6009490887 |
| Total numbers of transcripts | 162106 |
| Total transcriptomics length in bps | 113400385 |
| Average transcript length in bps | 699.54 |
| Maximum length of transcripts (>5000) bps | 331 |
| Minimum length of transcripts (200-300) bps | 77592 |
| N50 | 1356 |
| Total numbers of unigenes | 148871 |
| Total size of all unigenes in bps | 92863852 |
| Average unigenes length in bps | 623.78 |
| Maximum length of unigenes bps | 11998 |
| Minimum length of unigenes bps | 201 |
| N50 | 1167 |
| GC content | 40251760 (43.34%) |
| Depth of unigenes | 92863852 (65%) |
| Total number of CDS | 75198 |
| Total size of all CDS in bps | 49542030 |
| Average CDS length in bp | 662.81 |
| Maximum CDS length in bps | 9525 |
| Minimum CDS length in bps | 297 |
| Total annotated CDS hits with BLASTX | 71031 |
| Total without annotated CDS hits with BLASTX | 4167 |
| N50 | 789 |
| Unigenes in Uniprot databases | 42256 |
| Unigenes in COG database | 25444 |
| Unigenes in GO database | 6336 |
| Unigenes in Pfam database | 25809 |
| Unigenes with KEGG database | 6940 |

A total of 148871 unigenes, with a minimum length of 200–300 bp (highest unigenes), were generated, while 201 unigenes, with a maximum length of 900–1000 bp (minimum unigenes), were generated **(Supplementary Figure 2)**. GC [40251760 (43.34%)] and depth content [92863852 (65%)] of unigenes were obtained and are shown in **(Supplementary Figure 3).** Finally, 11998 unigenes were identified, with an average N50 length of 1167 bp (nucleotide), with a 623.78 average length. Amongst them, a total of 691 up-regulated and 100 down-regulated unigenes related to diosgenin metabolic biosynthesis pathways were annotated. **Table 1** show all transcriptome data collected from whole de novo sequencing and generated using a custom-made Perl script (Lulin et al., 2012; Nakasugi et al., 2013; Lehnert et al., 2014). The correlation indices between repeated samples wereD>D0.9, indicating that the Illumina sequencing results are credible. The raw reads of *Pedalium murex* were generated from Illumina *Hiseq* 2000 sequencing of all parts tissues and were submitted to the National Centre for Biotechnological Information (NCBI) Sequence Read Achieve (SRA) database with accession number-PRJNA787916, respectively.

### Functional annotation

The analysis of transcriptome data for functional characterization provides a global picture of biological processes, molecular and cellular activities, and the abundance of biosynthetic pathways. As a non-model plant, assembled transcripts of *P. murex* were aligned with six public protein databases to provide the best annotations. All assembled unigenes were searched against the Non-redundant (Nr), Uniprot, Kyoto Encyclopedia of Genes and Genomes (KEGG), Pfam, Gene Ontology (GO), and Clusters of Orthologous Groups (COG) databases using the BLASTx program with an E-value <1e-6. A total of 148871 unigenes were functionally assembled, in which 25444, 42256, 25809, and 6940 respective unigenes of COG UniProt, Pfam, and KEGG showed significant similarity to known proteins in NR databases **(Figure 2).** The result of BLASTX with different databases of clusters of orthologous groups of proteins (COG) and their annotation is listed in **(Figure 3)**. Genes of *P. murex* in an orthologous relationship were classified. Based on the Nr database, the E-value distribution indicated that 71031 of the matched unigenes ranged from 1e-6 to 1e-100 and showed the higher unigenes number of *Sesamum indicum,* while the higher similarity of these herbs with *Erythranthe guttata* and *Olea europaea var. sylvestris* is shown in **(Figure 4).** For the similarity distribution, a total of 148871 unigenes were exhibited.

**Figure 2:**
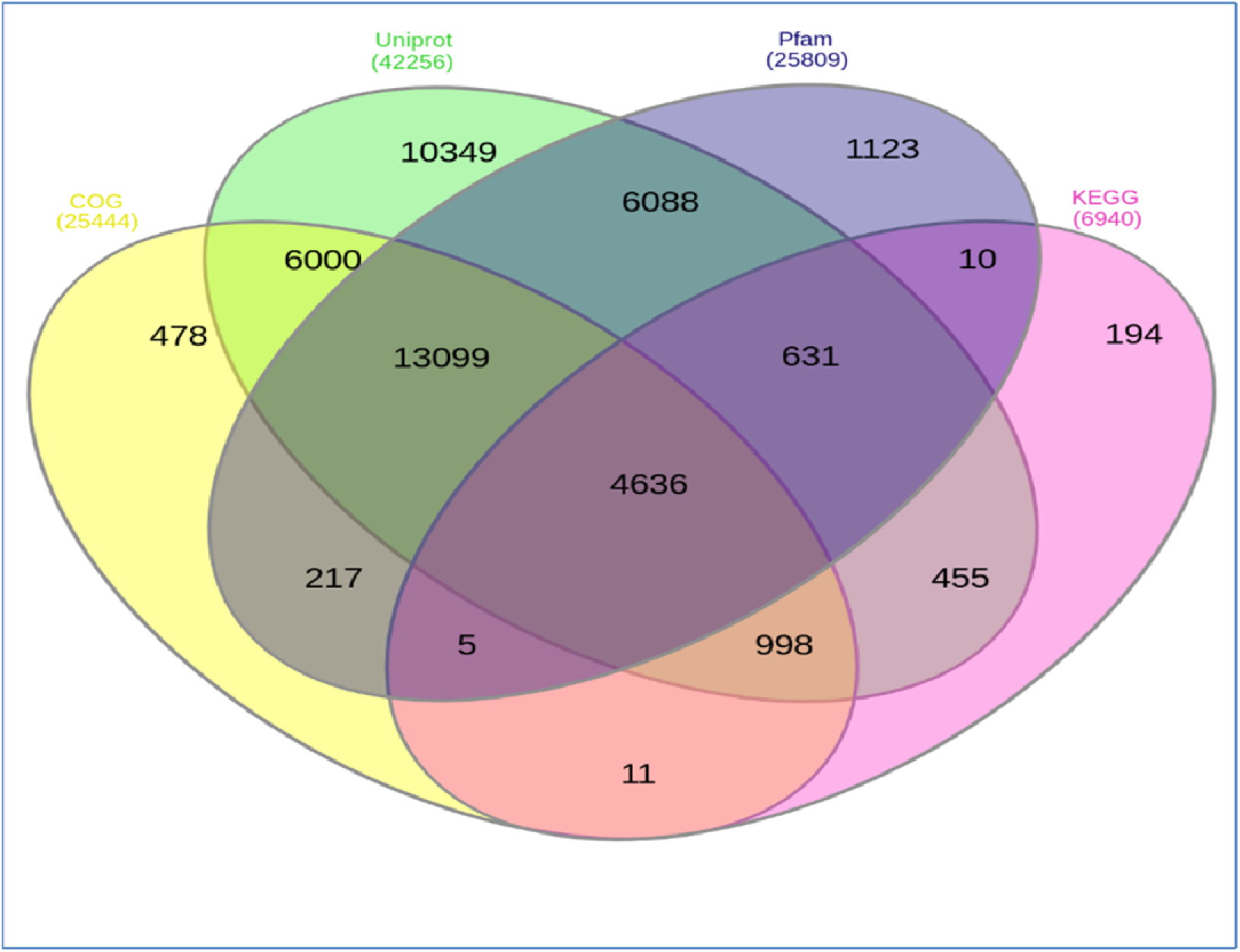
Functional annotation: Venn diagram showing differences common unigenes amongst them COG, Uniprot, Pfam, and KEGG databases of *P. murex*.

**Figure 3:**
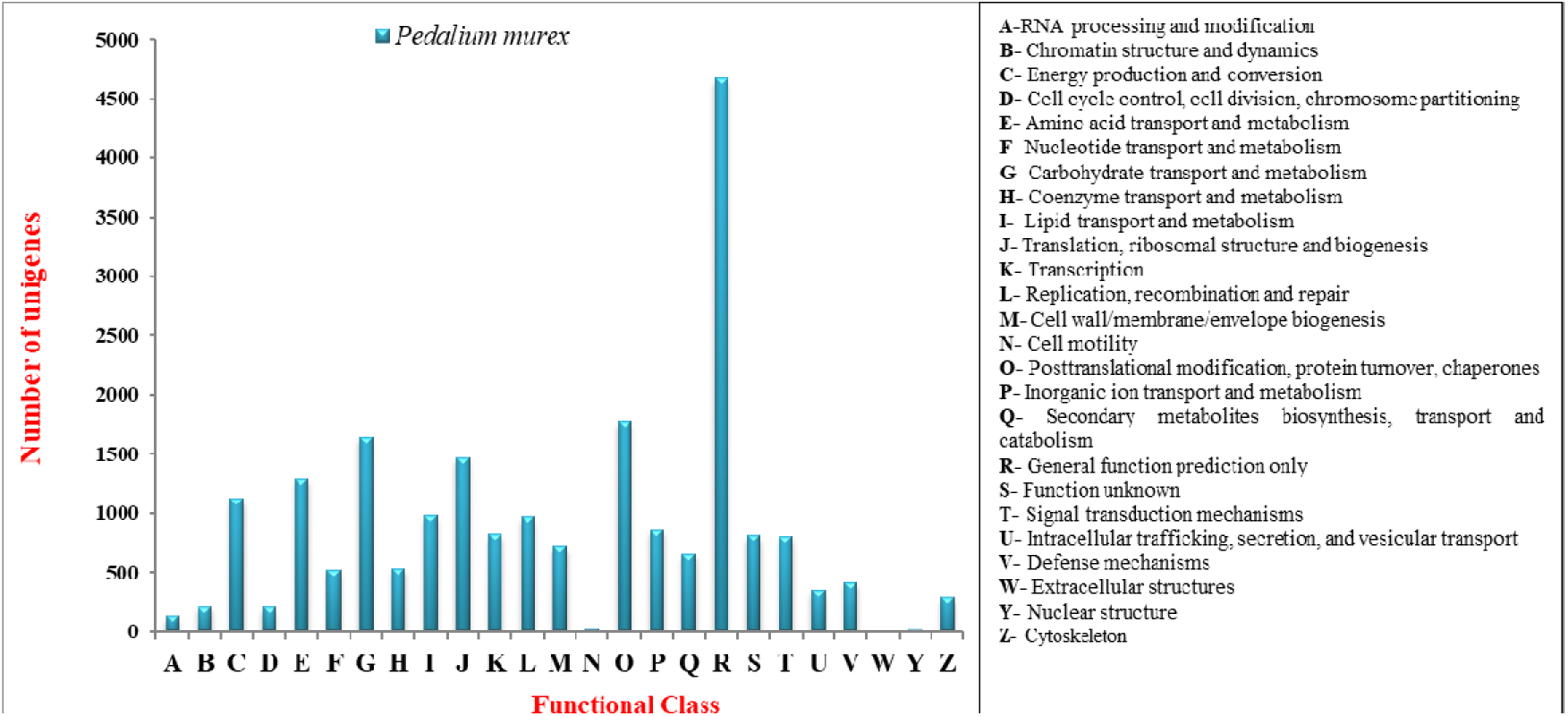
Functional annotation classifications: Cluster Orthologous Groups (COG) classification of transcripts into 25 categories; database of *P. murex*.

**Figure 4:**
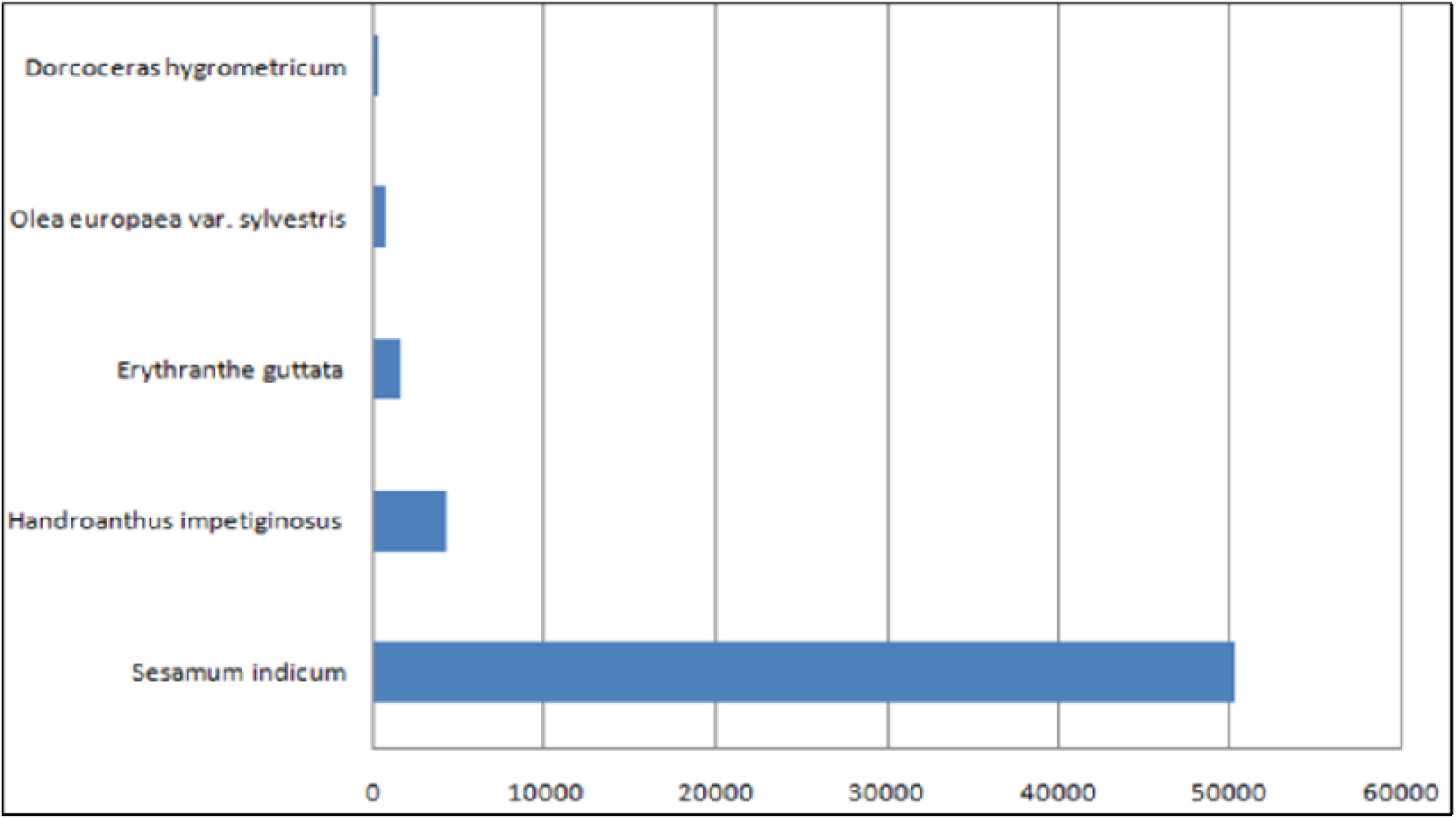
Top hit homologous Similarity of *P. murex*: Similarity of unigenes annotated using the Nr database. E-value distribution of best BLAST hits for each unigene (E-value <1e-6), similarity distribution of top BLAST hits for each unigenes. X-axis represents species and Y-axis represents number of unigenes.

For functional analysis and inferring the biological importance of genomic and transcriptome information, gene ontology (GO) has been widely used (Gene Ontology, 2004). GO analysis included three domains describing biological processes, cellular components, and molecular functions. When GO was used to classify gene functions, 13432 unigenes were assigned to 60 functional categories. Within the 4748 unigenes of the biological process (BP) domain, operations or sets of molecular events with a defined beginning and end are pertinent to the functioning of integrated living units: cells, tissues, organs, and organisms. In the 3418 unigenes of the cellular component (CC) domain, the parts of a cell or their extracellular environment are described. In the 5266 unigenes of the molecular function (MF) domain, the elemental activities of a gene product at the molecular level, such as binding or catalysis, are shown in **Figure 5 and Table 2**. To better understand the functions of specific metabolic pathways in *P. murex*, we mapped the annotated unigenes to the reference biological pathways in the KEGG database. A total of 75198 CDS were assembled and compared against the KEGG databases using BLASTX with a threshold bit-score value of 60 (default. A total of 309 pathways were found, of which 11 were metabolic, 4 were genetic information processing, 3 were environmental information processing, 5 were cellular processes, 10 were organismal systems, 12 were human diseases, and 3 were BRITE hierarchies’ pathways.

**Figure 5:**
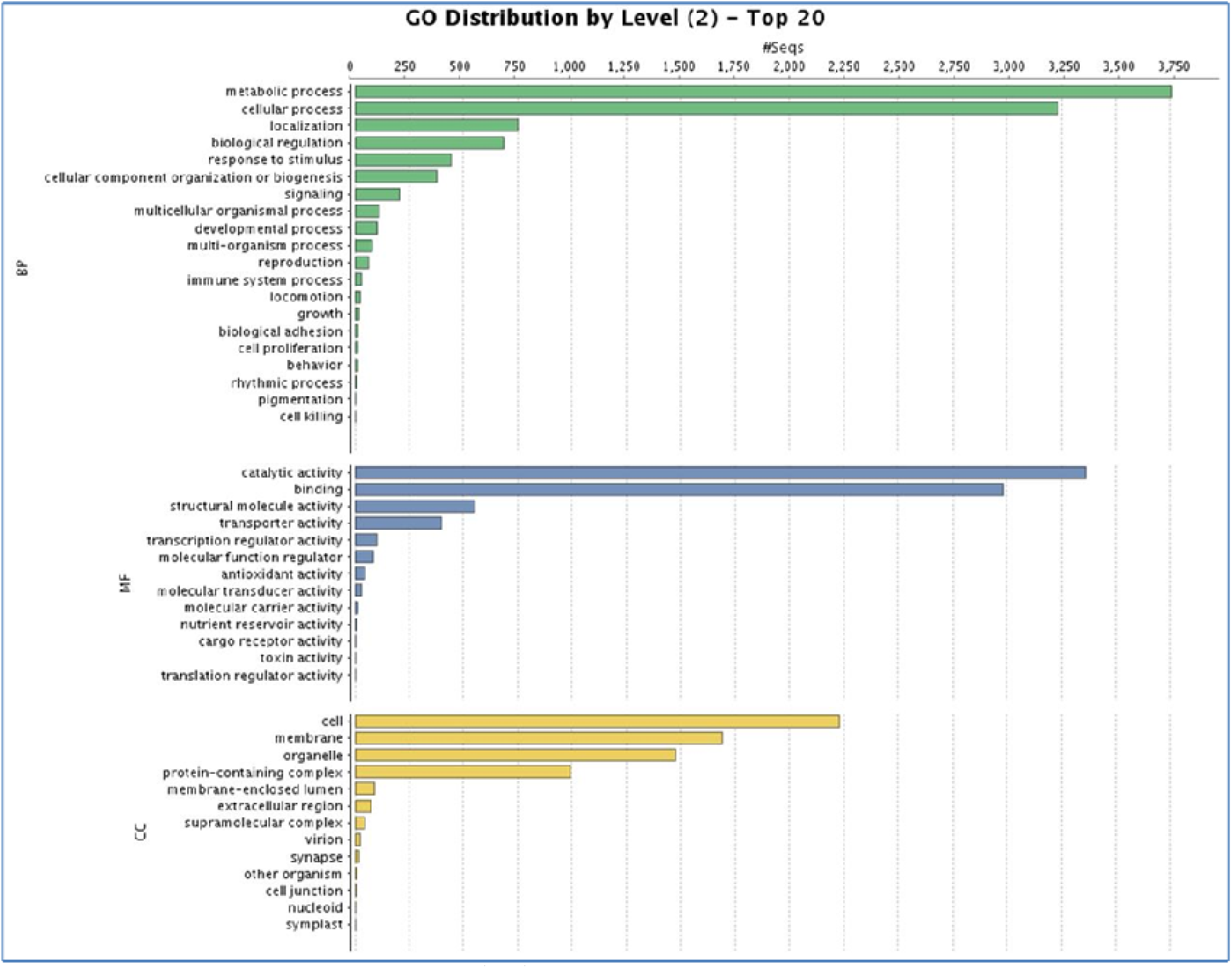
Comparative Gene Ontology (GO) classifications of commonly expressed functionally annotated differentially expressed genes (DEGs) from tissues of *P. murex* transcriptome. The genes corresponded to three main categories, cellular component (CC), molecular function (MF), and biological process (BP).

**Table 2:**
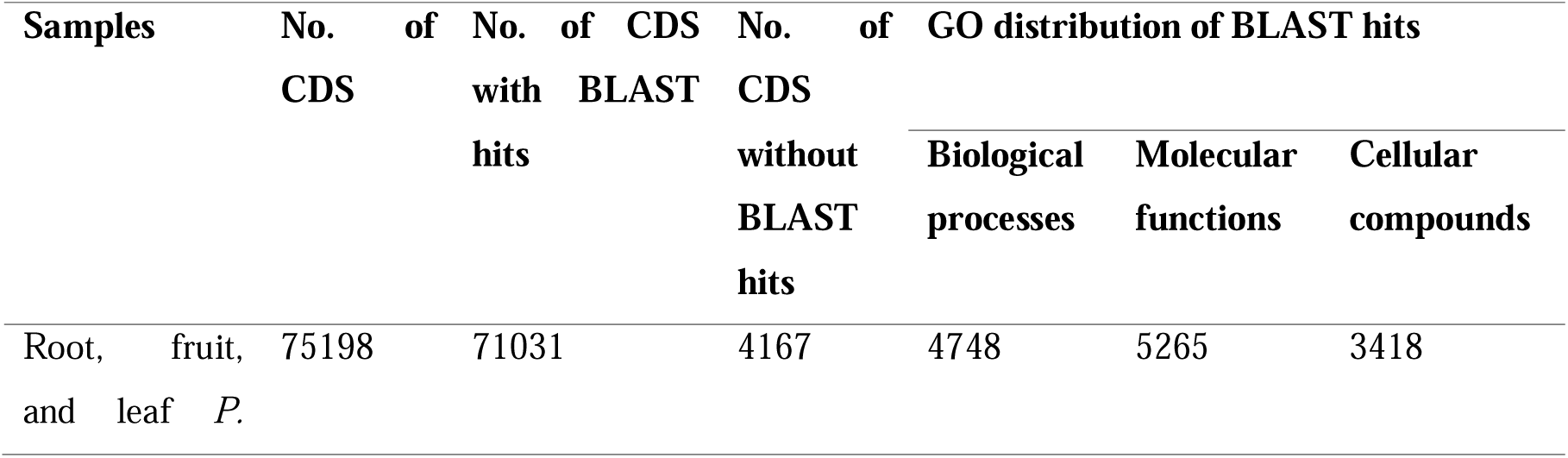
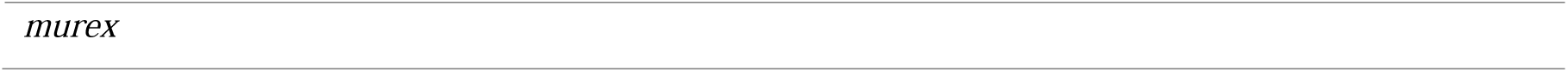
Transcriptome distribution of CDS with the BLASTX of all parts of *P. murex*.

Kanehisa et al. (2000) provide context for ongoing metabolic processes within an organism, allowing for a better understanding of the transcripts’ biological functions. A total of 75198 unigenes could be assigned to six main categories and 45 sub-categories, as shown in **Figure 6 and Table 3**. These enzymes feature assigned functions in 3140 CDS metabolic pathways in KEGG. Among these unigenes, 118 CDS encode key enzymes that are related to the pathways for terpenoid biosynthesis, including the synthesis of the terpenoid backbone (30 unigenes), monoterpenoids (3 unigenes), diterpenoids (20 unigenes), sesquiterpenoids and triterpenoids (7 unigenes), and other terpenoid-quinone complexes (43 unigenes). A total of 5 unigenes of tropane, piperidine, and pyridine and five unigenes of isoquinoline alkaloid biosynthesis are involved. Exactly 91 unigenes were associated with the flavonoid biosynthesis pathway, including the phenylpropanoid (45 unigenes), flavonoid (44 unigenes), flavone and flavonol (2 unigenes) biosynthesis pathways.

**Figure 6:**
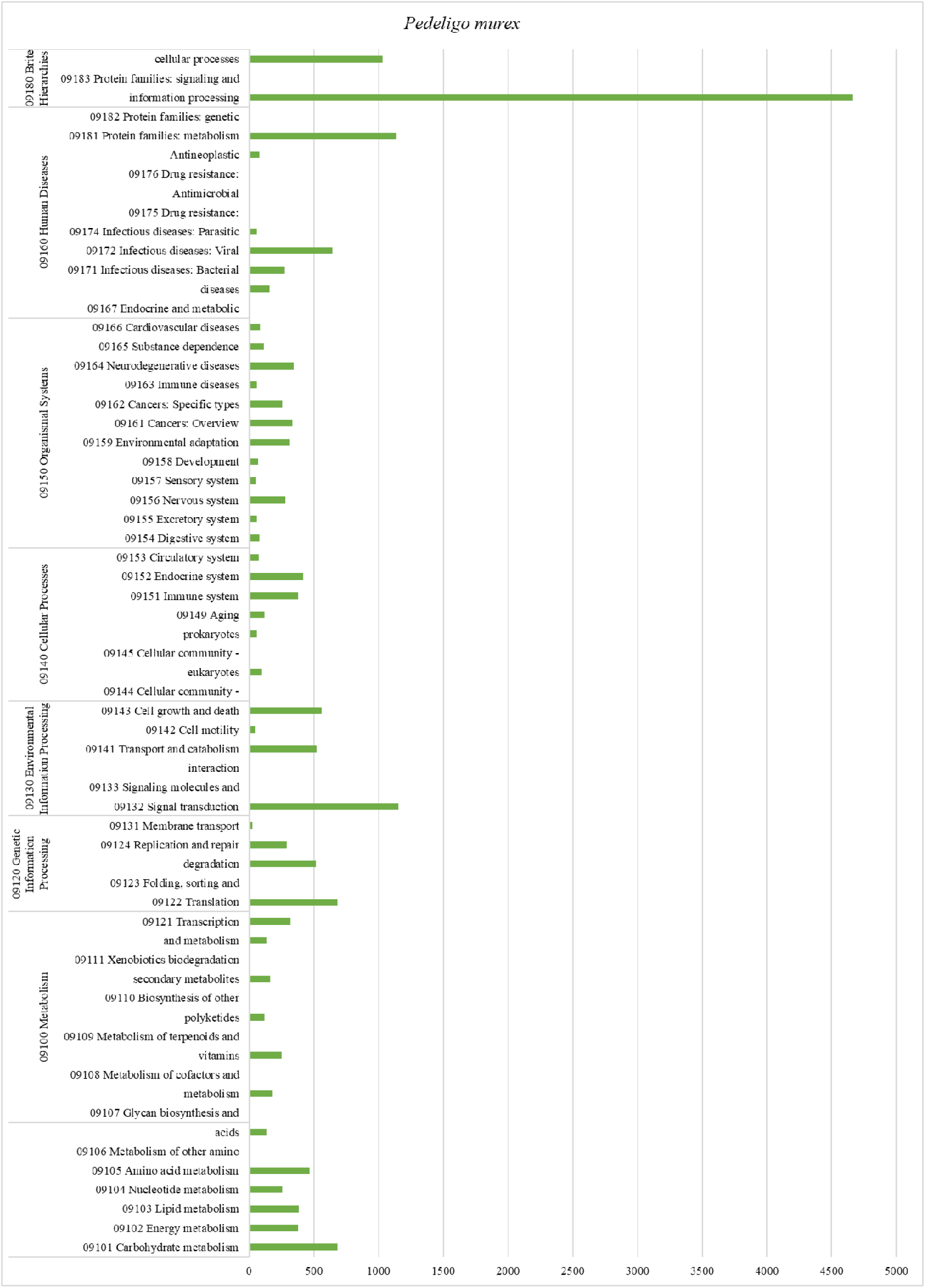
Functional annotation classifications: Kyoto Encyclopedia of Genes and Genomes (KEGG) classification details of six main categories, I: Metabolism, II: Genetic information processing, III: Environmental information processing, IV: Cellular processes, V: Organismal systems, (VI) Human diseases of *P. murex*. X-axis represents no. of CDSs and Y-axis represents KEGG functional pathway categories. X-axis represents no. of CDSs and Y-axis represents KEGG functional pathway categories.

**Table 3:**
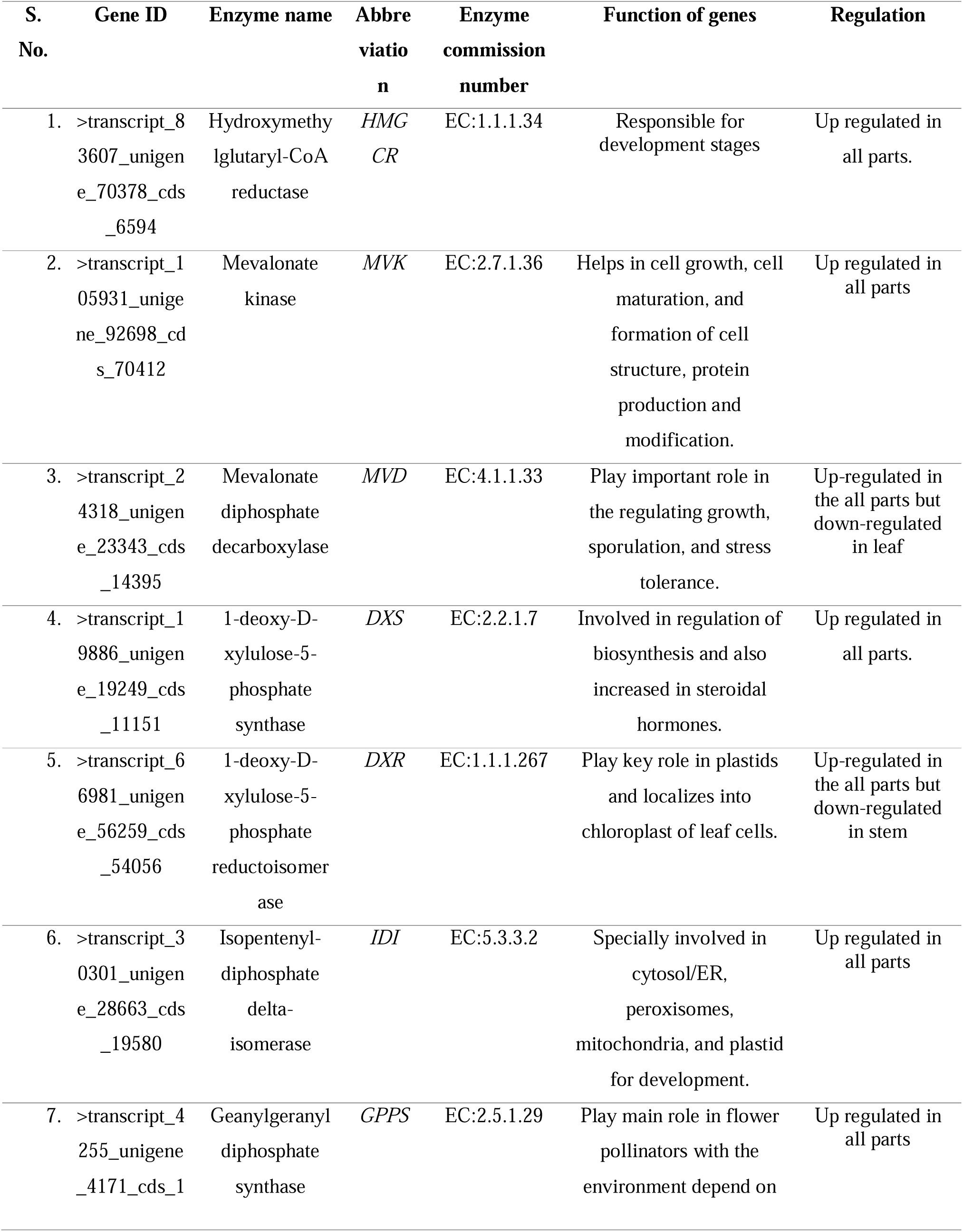
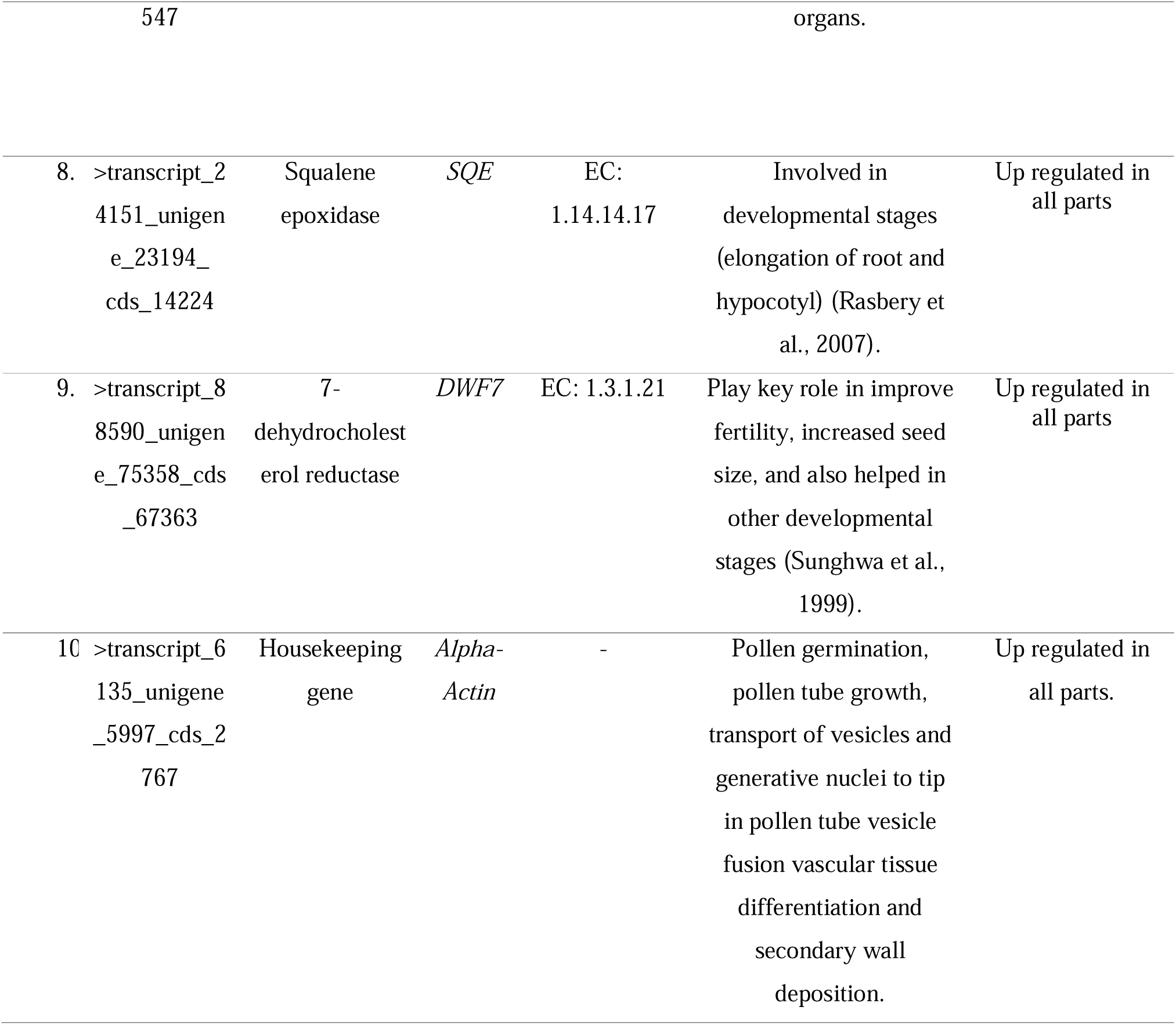
List of genes which are related to diosgenin in *P. murex*.

A total of 17 isoform were expressed in up-regulate in *P. murex* and these isoforms (unigene_23343_cds_14395; unigene_80426_cds_68315; unigene_23344_cds_14396; unigene_145953_cds_7445; unigene_28663_cds_19580; unigene_23095_cds_14117; unigene_4171_cds_1547; unigene_75358_cds_6736; unigene_56259_cds_54056; unigene_22026_cds_13237; unigene_22026_cds_13234; unigene_23194_cds_14224; unigene_22026_cds_13236; unigene_22026_cds_13235; unigene_70378_cds_65948; unigene_19249_cds_1115) were assembled in the diosgenin pathway analysis.

Meanwhile, the top 20 significant pathways were analyzed based on the FDRD≤D0.05. *P. murex* root, fruit, and leaf tissues in “biological process” metabolic process, cellular process, localization, biological regulation, response to stimulus, cellular component organization or biogenesis, signalling, multicellular organismal process, developmental process, multi-organism process, reproduction, immune system process, locomotion, growth, biological adhesion, cell proliferation, behaviour, rhythmic process, pigmentation, cell killing, diosgenin biosynthesis, and photosynthesis, terpenoid backbone biosynthesis; in “molecular function” catalytic activity, binding, structural molecular activity, transporter activity, transcription regulator activity, molecular function regulation, antioxidant activity, molecular carrier activity, nutrient reservoir activity, cargo receptor activity, toxin activity, transcription regulator activity; in “cellular process” cell, membrane, organelle, protein-containing complex, membrane-enclosed lumen, extracellular region, super molecular complex, virion, synapse, other organism, cell junction, nucleoid, symplast biosynthesis showed significant enrichment are shown in **(Figure 7) and Table 2** In addition to the common pathways of primary metabolism, enriched secondary metabolic pathways, including terpenoid and diosgenin biosynthesis, were also found between different tissues of *P. murex*, indicating the possible distinct distribution of secondary metabolites in this herb.

**Figure 7:**
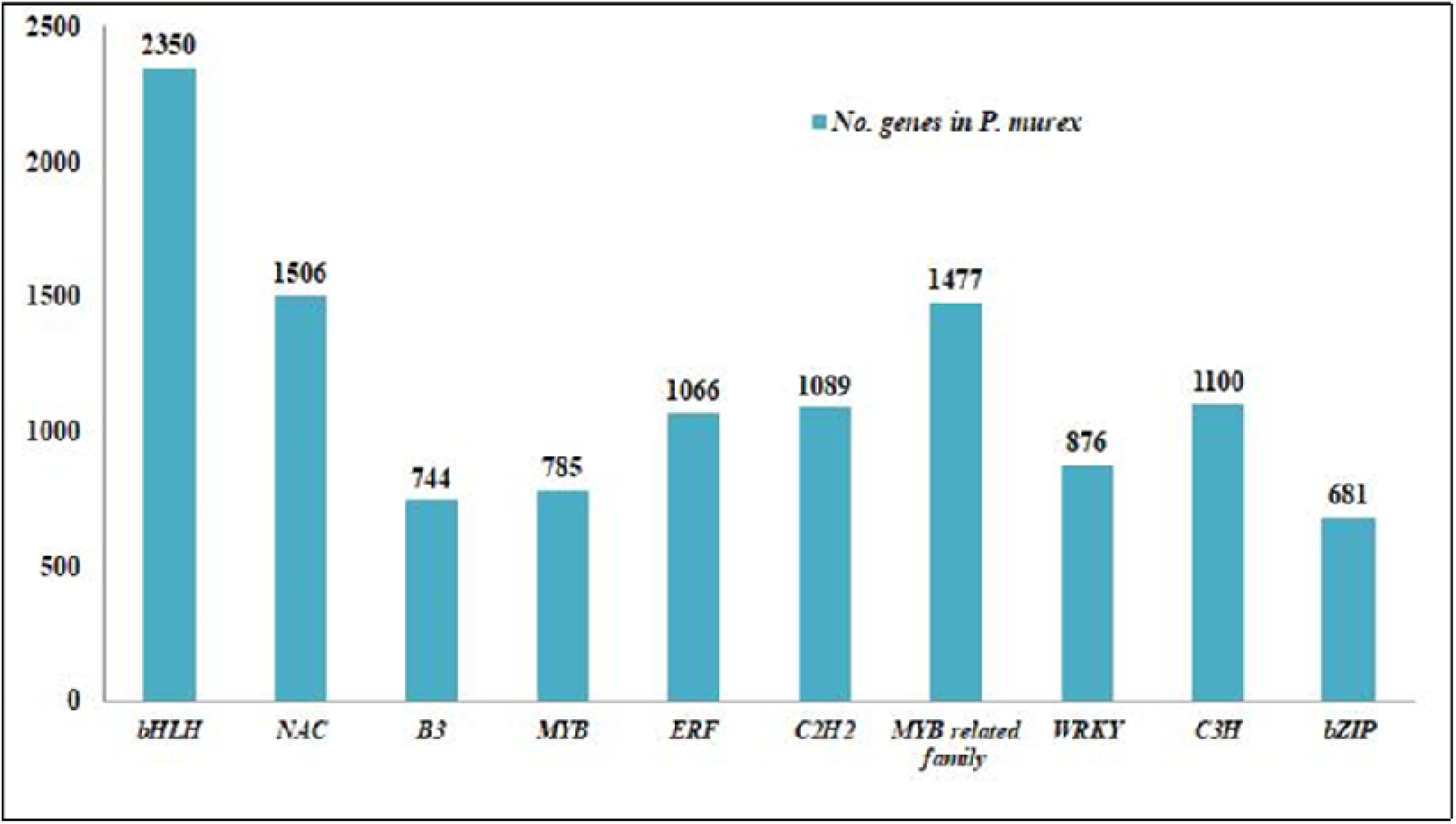
Functional annotation classifications: Classification of transcripts into major transcription factor (TF) families of *P. murex*.

### Identification of Transcription Factors

Transcription factors (TFs) attach to specific cis-regulatory components of promoter regions and play an essential role in gene expression, plant secondary metabolism, and plant response to environmental stress. In numerous plant species, TF families such as ARF, bHLH, bZIP, MYB, NAC, and WRKY have been implicated in secondary metabolite regulation as well as abiotic and biotic stress responses (Fujita et al., 2006; Patra et al., 2013). A total of 21026 unigenes of transcription factors were annotated and were grouped into ten transcription factor families, namely, bHLH (2350), NAC (1506), MYB related family (1477), ERF/AP2 (1066), C2H2 (1089), WRKY (876), C3H (1100), bZIP (681), MYB (785), B3 (744) were highly expressed. Members of the WRKY, bHLH, and AP2/ERF families have been shown to have significant roles in regulating the biosynthesis of terpenoids, alkaloids, and steroids, as shown in **Figure 7**. The most abundant were members of the bHLH, MYB-related family, and NAC. They constitute a valuable gene resource for further studies of their regulatory functions in diosgenin biosynthesis.

### qRT-PCR validation of selected genes

To verify the expression profiles obtained from Illumina sequencing, we performed qRT-PCR on nine selected genes related to triterpene diosgenin biosynthesis **Figure 9**. In this experiment, each qRT-PCR reaction contained 1 µl of diluted cDNA of root, fruit, and leaf, 1 µl of each primer, 5 µl of 2X SYBR Green mix, and 2 µl of RNase-free water. All qRT-PCRs were performed using the following conditions: denaturation at 95 °C for 1 min, followed by 40 cycles of 95 °C for 20 s and then at 60 °C for 1 min. Successive qRT-PCR assays were performed using triplicates, and to verify product specificity, melting curve analysis was performed after each amplification. The alpha-tubulin gene of *P. murex* was used as an internal reference gene for the normalization, and three technical replicates were performed for each sample. Relative expression levels for each gene were calculated using the 2−ΔΔCt method (Livak et al., 2001).

**Figure 9:**
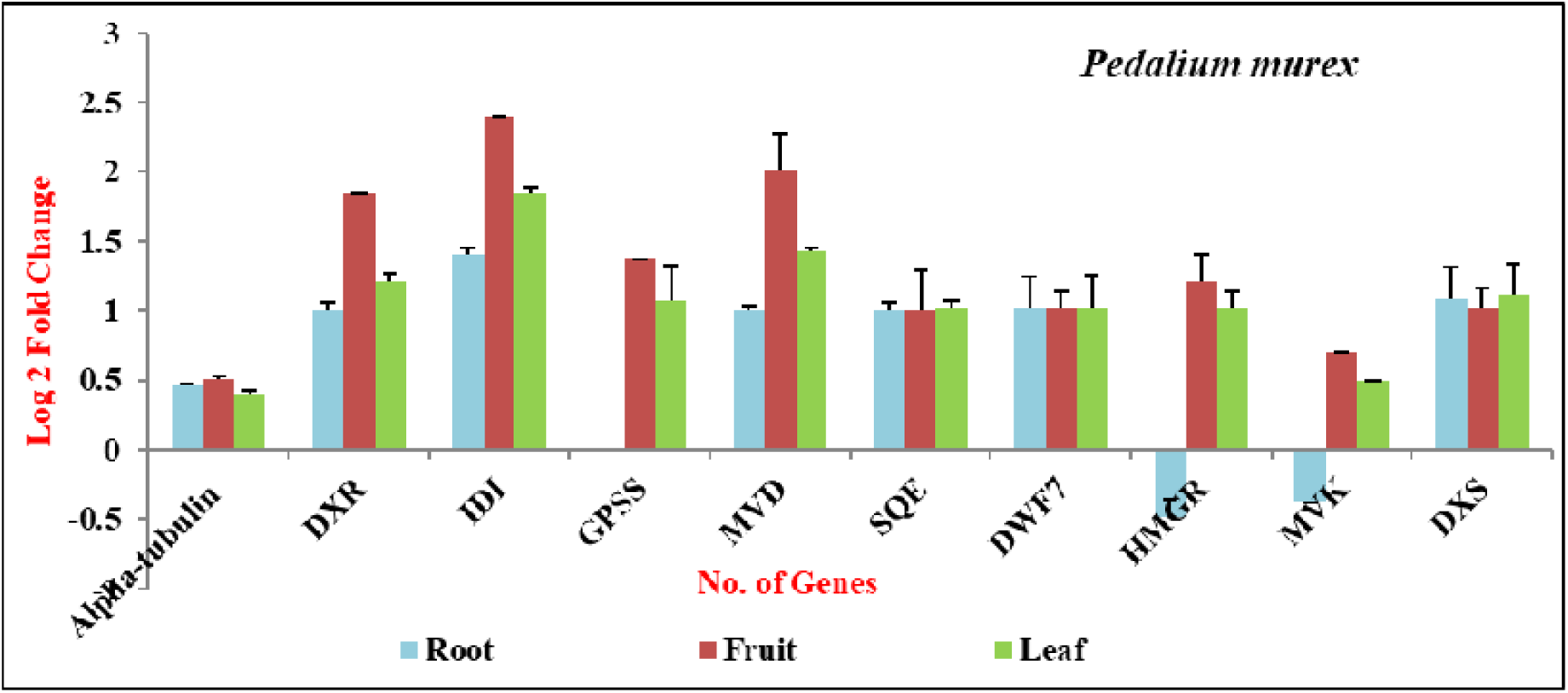
qRT-PCR validation of differential 9 selected genes related to diosgenin biosynthesis of *P. murex*. The relative transcript levels were normalized with *Alpha tubulin* as the standard. *HMGCR*: hydroxyl methylglutarylCoA reductase; *MVK*: mevalonate kinase; *MVD*: Mevalonate diphosphate decarboxylase; *DXS*: 1-deoxy-D-xylulose-5-phosphate synthase; *DXR*: 1-deoxy-D-xylulose-5-phosphate reductoisomerase; *IDI*: Isopentenyl-diphosphate delta-isomerase; *GPPS*: Geranyl diphosphate synthase; *SQE*: Squalene epoxidase. X-axis represents no. of genes and Y-axis represents number of Log 2-Fold change.

Consistent with the Illumina data, most of these genes showed intense expression levels in root, fruit, and leaf tissues of *P. murex,* and acetyl-CoA acetyltransferase, hydroxymethylglutaryl-CoA synthase, hydroxymethylglutaryl-CoA reductase, and squalene epoxidase (*SQE*) genes were expressed abundantly. The expression fold changes were also close to the RNA-*seq* results. The qRT-PCR results show that the RNA-*seq* data in this study was accurate. A total of 10 genes were identified in the transcriptome data analysis of the root, fruit, and leaves of *P. murex,* which are related to diosgenin biosynthesis. Of those, nine genes were selected for validation by Quantitative Real-Time Polymerase Chain Reaction (qRT-PCR). The primers for these selected genes were used in the qRT-PCR analysis and are listed in **Table 3**.

We can use metabolic pathway analysis to learn about the relationships between genes in each pathway and their biological activities. Triterpenes are synthesized from a five-carbon isoprene unit through the cytosolic MVA pathway. Diosgenin is composed of six isoprene units derived from the C-30 hydrocarbon precursor, squalene. Squalene is synthesized from isopentenyl diphosphate *(IPP)* via the MVA pathway. All genes encoding the enzymes associated with the up-and-down-regulate regions of steroid diosgenin biosynthesis were successfully identified in the *P. murex* transcriptome. Their expression values were monitored in three biological replicates, along with their mean values. In particular, the Hydroxymethylglutaryl-CoA reductase (*HDMGR)* gene in leaf, geranylgeranyl diphosphate synthase (*GPPS)* gene in root, and Isopentenyl-diphosphate delta-isomerase *(IDI)* gene in fruit are highly expressed, in which case the 1-deoxy-D-xylulose-5-phosphate reductoisomerase *(DXR)* gene in fruit, *IDI* gene in fruit, *GPPS* gene in fruit, and mevalonate kinase *(MVK)* gene in leaf, fruit, and root are not expressed compared to others are shown in **Figure 7, and Table 4**. Most unigenes related to the MVA pathway were specifically up-regulated in the leaf and root of the MEP/DOX pathway of *P. murex*. In **Additional File 2,** list of transcripts related to triterpenoid diosgenin backbone biosynthesis is shown.

**Table 4:**
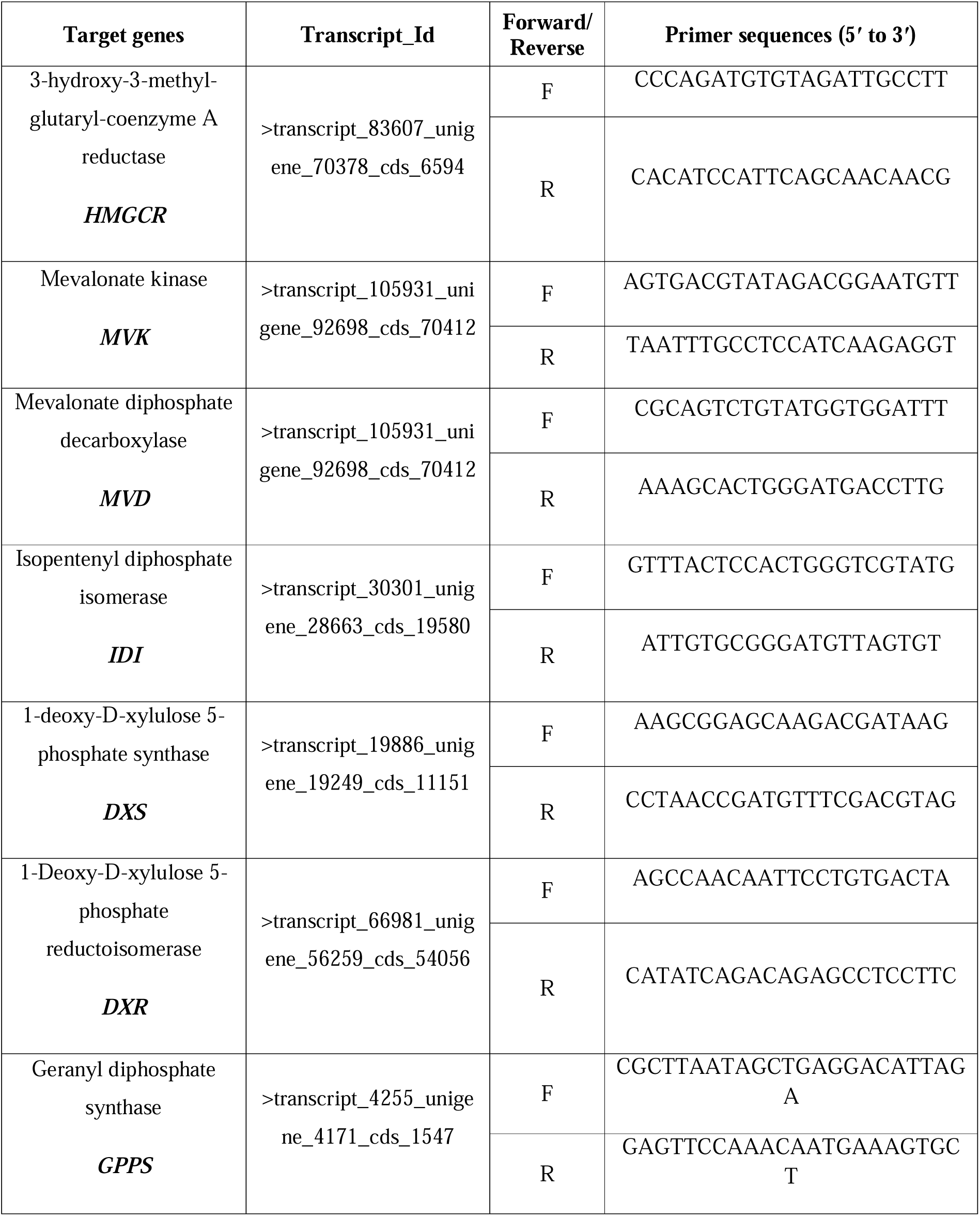
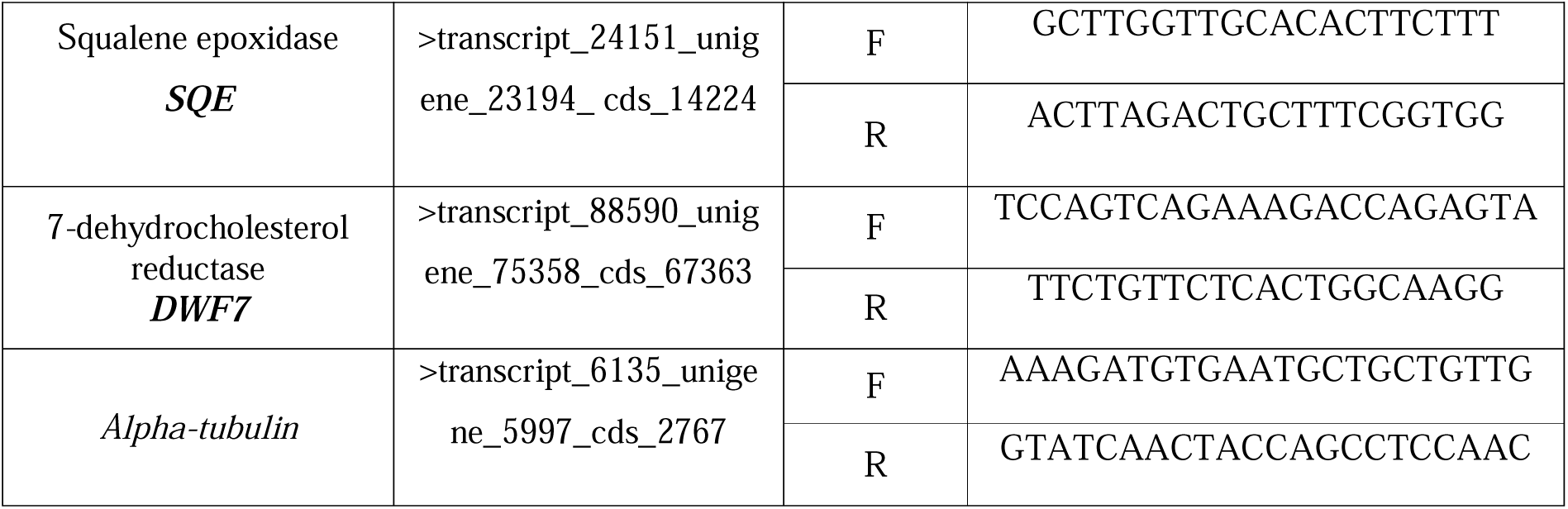
Primer Identification of internal control genes and some of these genes for validation of qRT-PCR analysis of *T. terrestris* and *P. murex*.

### Identification of SSRs

The most important molecular markers are simple sequence repeats (SSR) and microsatellites, used for gene mapping, molecular breeding, and genetic diversity. These are the tandem repeats of nucleotide motifs of sizes 2-6 bp, and they are highly polymorphic and are ubiquitously present in all the known genomes. Thus, SSRs were identified from assembled transcript sequences with the MISA perl. SSRs having a flanking of 150 bp (upstream as well as downstream) were fetched with an in-house python script, which can be used for primer designing. A total number of 8760 SSRs loci with di-, tri-, tetra-, penta-, and hexanucleotide repeats were recognized, in which 6108 of di-nucleotide, 2502 of trinucleotide, 146 of tetra-nucleotide, and 4 of penta-nucleotide motifs were found **(Figure 10 and Table 5)**.

**Figure 10:**
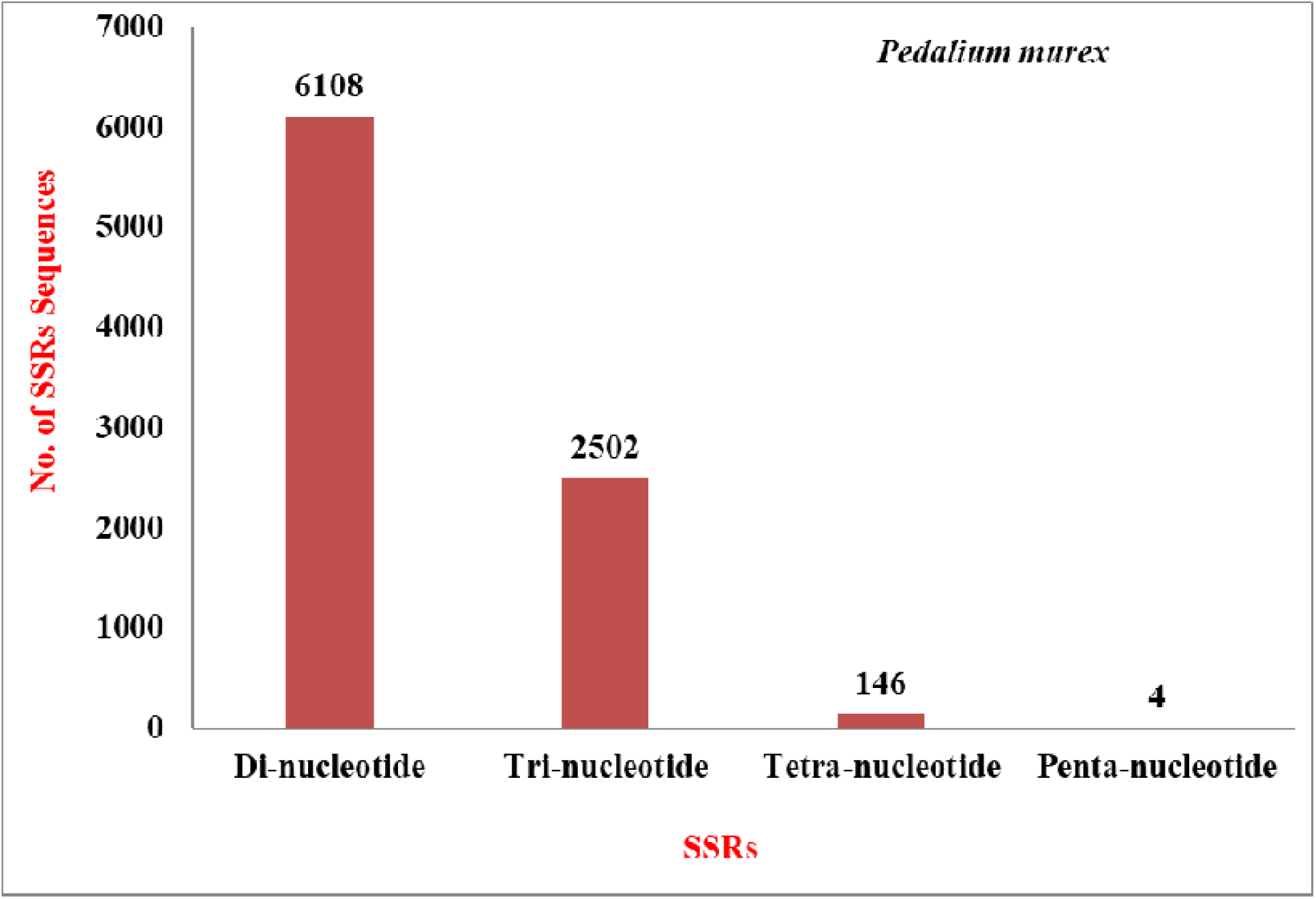
Graph 24: Frequency of the individual repeat nucleotides in the SSRs obtained from MISA analysis of *P. murex*. X-axis represents SSRs repeats, and Y-axis represents number of assembled sequences.

**Table 5:** Statistical analysis of SSR’s identification from *P. murex*.

| SSR data of all parts of *P. murex* | Number |
| --- | --- |
| Total number of sequences examined | 162106 |
| Total size of examined sequences (nt) | 113400385 |
| Total number of identified SSRs | 8760 |
| Numbers of SSR containing sequences | 7749 |
| Numbers of sequences containing more than 1 SSR | 881 |
| Number of SSRs present in compound formation | 517 |
| Di-nucleotide | 6108 |
| Tri-nucleotide | 2502 |
| Tetra-nucleotide | 146 |
| Penta-nucleotide | 4 |

## Discussion

Medicinal plants produce a significant number of pharmaceutically and industrially relevant secondary metabolites via complex metabolic processes. *Pedalium murex* is an important traditional medicinal *herb* with numerous pharmaceutical activities. Both herbs are pharmacologically significant, but there needs to be more research on genomics and the transcriptome of *P. murex*. However, the introduction of next-generation sequencing (NGS)-based high-throughput transcriptome sequencing (Metzker, 2010; Claros et al., 2012) has aided in overcoming the obstacles in such plants. In several non-model plants, the NGS technique has successfully elucidated important genes and regulators of complex metabolic networks. The NGS technique has also been implicated in the elucidation of molecular mechanisms of complicated biosynthetic processes using a *de novo* spatial transcriptome sequencing strategy in *T. govanianum,* as done in this study (Gahlan et al., 2012; Chakrabarty et al., 2015; Jayaswall et al., 2016). We performed a comparative transcriptome analysis of the root, fruit, and leaf tissues of *P. murex*. The dataset reported here is useful in understanding the biosynthetic pathway of diosgenin.

A total of ∼6.77 GB of clean data was generated for the root, fruit, and leaf of *P. murex*. Sample-wise paired-end reads of this herb (410 million) collected in this work were sufficient for reliable *de novo* transcriptome characterization and precise gene expression pattern quantification. 16889795 (41.10%) duplicate filtered reads assembled into 148871 unigenes with an average N50 length of 1167 bp, which is like that of previously reported non-model herbs and plants, *Trigonella foenum-graecum*, *Trillium govanianum,* and *Dioscorea zingiberensis* (Chen Zhou et al., 2019). GC content (65%) may be attributed to P. murex’s ability to adapt to extreme temperatures because GC content is important in gene regulation, physical description of the genome, and nucleic acid stability (Smarda Petr et al., 2014), *P. murex* (65%) had a GC content that was higher than *Arabidopsis* (42.5%) (Devi et al., 2016). Despite being a non-model plant, the annotation of *P. murex* transcripts with multiple public databases successfully assigned putative functions to over 7103 of the transcripts. Nonetheless, 4167 transcripts could not be annotated, possibly belonging to the un-translated regions, or representing the species-specific gene pool (Jung-A Kim et al., 2019). The best match for each unigenes search against the Nr and KEGG databases was of help to assign GO functional annotation under biological process, cellular component, and molecular function categories.

The presence of multiple gene families in *P. murex* is suggested by the assignment of GO keywords to a significant number of transcripts. The mapping of transcripts with the KEGG database in this work identified all the genes connected to the steroidal diosgenin pathway, which aids in understanding the biological function and interaction of genes related to primary and secondary metabolites. A total of 25444 transcripts were annotated and grouped into 25 functional categories using the KOG classification system. The importance of transcription factors (TFs) in modulating gene expression by binding to the promoters of single or many genes has long been recognized. Our results showed that transcription factor families (bHLH, NAC, MYB related family, ERF, C2H2, WRKY, C3H, bZIP, MYB, and B3) that help regulate secondary metabolites in plants were substantially represented as shown in **(Figure 7)**. TSAR1 (Triterpene Diosgenin Biosynthesis Activating Regulator1) and TSAR2 are members of the bHLH family and regulate triterpene diosgenin biosynthesis in *Medicago truncatula* (Goossens et al., 2015). As a result, the identified TFs in this study can be investigated as potential regulators of steroidal diosgenin biosynthesis in *P. murex*.

On the other hand, RNA-*seq* is unable to discover genes or transcripts with low expression levels, which are particularly relevant for regulatory genes (Jiang et al., 2015; Malone et al., 2011). The transcription factors (TFs) involved in sexual dimorphism in flies, doublesex (*dsx*) and fruitless (*fru*), cannot be detected by RNA-*Seq* in the fully sequenced modENCODE embryo samples with low expression (Graveley et al., 2011). These genes will influence both estimates of how transcripts are expressed and research into how they work.

Nowadays, gene expression analyses are extremely widely utilized in the identification of putative regulators of complex molecular pathways by measuring transcriptional levels in different tissues and developmental stages in numerous fields (Loven et al., 2012). Glycolytic, mevalonate (MVA), MEP/DOX, and steroidal pathways are involved in diosgenin biosynthetic pathway. In our transcriptomic analysis, sequences encoding *HMGCR* (EC: 1.1.1.34)*, PMK* (EC: 2.7.4.2)*, IDI* (EC: 5.3.3.2), and *MVD* (EC: 4.1.1.33) (a rate limiting enzyme) genes of *P. murex* represented the highest number (1, 2, and 2 respectively) of unigenes associated with the MVA pathway. This pathway, involved in the synthesis of phytosterols, carotenoids, gibberellins, triterpenoid and steroidal saponins, was found highly expressed in fruit followed by leaf and least expression in root, therefore suggesting its possible role in the early stages of fruit development and defence by producing derivatives of saponins (Kim et al., 2014). *P. murex DXS* (EC: 2.2.1.7) and *DXR* (EC: 1.1.1.267) genes encode the highest number (1 and 1, respectively) of unigenes associated with the MEP/DOX pathway. *GPPS* (EC: 2.5.1.1) and *SQE* (EC: 1.14.13.132) genes of *P. murex* with the associated squalene biosynthesis, in which the *GPPS* gene represented the highest number (6 transcripts) **(Table 5).** Higher expression of MEP pathway genes in aerial parts is in accordance with earlier studies (Devi et al., 2016).

The cyclization of 2, 3-oxidosqualene is a branch point of diosgenin synthesis. The major diosgenin related genes are identified in the fruit and leaf of *P. murex*. Among the three transcripts of *SMT1* (EC: 2.1.1.41) and six transcripts of *DWF7* (EC: 1.3.1.21), 3 were identified in the steroidal biosynthesis pathway. A total of 6 transcripts were annotated as housekeeping genes (alpha-tubulin) in our transcriptome. The highest expression of *alpha-tubulin* synthase in the fruit of *P. murex* makes it an important enzyme related to triterpenoid diosgenin biosynthesis at later stages. Similar tissue-specific concentrations of triterpenoid diosgenin have already been reported in other herbs (Wei Sun et al., 2017; Chen Zhou et al., 2019). Moreover, *alpha-tubulin*, *HMGCR, DXR, IDI, GPPS, MVD, SQE, DWF7, HMGCR,* and *DXR* genes were up-regulated while the *HMGCR, MVK* gene in root was down-regulated in *P. murex.* Further characterization of these candidate enzymes is needed to confirm the pathway of triterpenoid diosgenin biosynthesis in *P. murex*.

The most important molecular markers are simple sequence repeats (SSR) and microsatellites, which are used for gene mapping, molecular breeding, and genetic diversity. These are the tandem repeats of nucleotide motifs of sizes 2-6 bp, and they are highly polymorphic and are ubiquitously present in all the known genomes. Thus, SSRs were identified from assembled transcript sequences with the MISA perl. SSRs having a flanking of 150 bp (upstream as well as downstream) were fetched with an in-house python script, which can be used for primer designing. In this research work, we found that total number of 15551 and 8760 microsatellites from *P. murex*, respectively have steroidal saponins properties. Amongst them, the identified SSRs, with di-, tri-, tetra-, penta- and hexanucleotide repeats were recognized in which, 9292 and 6108 of di-nucleotide, 6580 and 2502 of trinucleotide, 146 and 146 of tetra-nucleotide, 4 and 4 of penta-nucleotide motifs were found as shown in **Table 6 and Figure 10** in which, di-/trinucleotides are highly predominant while, pentanucleotide motif are lesser in *P. murex*.

## Conclusions

Pharmaceutical industries rely on medicinal plants as sources of botanical raw pharmaceuticals. The whole comparative transcriptome analysis of root, fruit, and leaf tissues of *P. murex* was carried out in this study. The experiment was carried out on *P. murex* to see if the potential genes involved in the biosynthesis of steroids and diosgenin can be examined in the future for up-scaling of the targeted bioactive compounds at the industrial scale of an important medicinal plant. In *P. murex,* the highest expression of critical pathway genes and regulatory candidates is seen in the root and leaf, which suggests they could be the sites of steroidal diosgenin production. The current study’s findings will serve as a foundation for multi-omics investigations in *P. murex* and similar species to better understand steroidal diosgenin production and accumulation. The information gathered will help find new genes and learn more about how *P. murex* works, which will help improve its genes and help protect it.

## Acknowledgement

We are thankful to the Director and Head of Department of Botany, Dayalbagh Educational Institute, Dayalbagh (Agra).

## Abbreviations

HMGCR: 3-Hydroxy-3-Methylglutaryl-CoA Reductase
MVK: Mevalonate Kinase
MVD: mevalonate diphosphate decarboxylase
IDI: Isopentenyl pyrophosphate isomerase
DXS: 1-Deoxy-D-xylulose-5-phosphate synthase
DXR: 1-deoxy-D-xylulose 5-phosphate reductoisomerase
GPPS: Geranyl diphosphate synthase
SQE: Squalene monooxygenase
DWF7: 7-dehydrocholesterol reductase
HMGS: Hydroxymethylglutaryl-CoA synthase
SQS: Squalene synthase
SMT1: Sterol methyltransferase 1
CYP51: Cytochrome P450
FK: Fructokinase
qRT-PCR: Quantitative Real-Time Polymerase Chain Reaction

